# The dorsal blastopore lip is a source of signals inducing PCP in the *Xenopus* neural plate

**DOI:** 10.1101/2021.04.26.441483

**Authors:** Pamela Mancini, Olga Ossipova, Sergei Y. Sokol

## Abstract

Coordinated polarization of cells in the tissue plane, known as planar cell polarity (PCP), is associated with a signaling pathway critical for the control of morphogenetic processes. Although the segregation of PCP components to opposite cell borders is believed to play a critical role in this pathway, whether PCP derives from egg polarity or preexistent long-range gradient, or forms in response to a localized cue remains a challenging question. Here we investigate the *Xenopus* neural plate, a tissue that has been previously shown to exhibit PCP. By imaging Vangl2 and Prickle3, we show that PCP is progressively acquired in the neural plate and requires a signal from the posterior region of the embryo. Tissue transplantations indicated that PCP is triggered in the neural plate by a planar cue from the dorsal blastopore lip. The PCP cue did not depend on the orientation of the graft and was distinct from neural inducers. These observations suggest that neuroectodermal PCP is not instructed by a preexisting molecular gradient, but induced by a signal from the dorsal blastopore lip.

**Highlights:**

- The *Xenopus* neural plate progressively acquires PCP in a posterior-to-anterior direction.
- The dorsal blastopore lip is likely the source of the PCP-instructing signal for the *Xenopus* neural plate.
- The PCP cue is distinct from neural inducers and has a planar mode of transmission.

## Introduction

During morphogenesis, collective cell behaviors may be orchestrated in a multicellular organism through the integration of individual polarities of constituent cells. The coordinated polarization of neighboring cells in the plane of the tissue is known as planar cell polarity (PCP). Genetic studies have implicated PCP components in many morphogenetic processes, including vertebrate neurulation, lung and kidney development and left-right patterning, and mutations in PCP genes have been linked to birth defects in humans (Goodrich and Strutt, 2011; Gray et al., 2011; Nikolopoulou et al., 2017; Tian et al., 2020). The core PCP pathway, initially identified in *Drosophila*, comprises two protein complexes: Frizzled/Dishevelled and Prickle/Van Gogh (Vangl in vertebrates) that segregate to opposite cell borders (Devenport, 2014; Goodrich and Strutt, 2011; Vladar et al., 2009). Both PCP complexes also include the atypical cadherin Flamingo (Celsr in vertebrates). These core proteins are part of a signaling pathway that regulates cell shape and motility through the action of multiple effectors, including components of the vesicular trafficking machinery and cytoskeleton-remodeling factors (Butler and Wallingford, 2017; Devenport, 2014).

PCP protein segregation to opposite cell sides is reinforced by reciprocal intracellular repulsion and extracellular stabilization of core complexes (Aw and Devenport, 2017; Fisher and Strutt, 2019; Peng and Axelrod, 2012), but an initial cue is needed to define the orientation of the polarity vector relative to the body axes. Molecular gradients have been considered as primary candidates for PCP-instructing signals (Chu and Sokol, 2016; Gao et al., 2011; Lawrence, 1966; Wu et al., 2013). Wnt protein gradients may instruct PCP by modulating Frizzled activity in the fly wing (Wu et al., 2013), or Vangl2 phosphorylation in the mouse limb bud (Gao et al., 2011), although the involvement of the fly Wnt ligands in PCP remains controversial (Ewen-Campen et al., 2020). Alternative candidate long-range PCP cues are mechanical strains (Aigouy et al., 2010; Aw et al., 2016; Chien et al., 2015). Studies of the *Xenopus* and mammalian epidermis suggested that physical forces generated during morphogenetic processes might act to define the global PCP axis (Aw et al., 2016; Chien et al., 2015). Nevertheless, the endogenous source of PCP cues and the mechanisms through which PCP is instructed at the tissue level remain to be fully elucidated.

Here, we use the *Xenopus* embryo to investigate the dynamics and the origin of PCP in the neuroectoderm, a tissue that is planar polarized in several vertebrate models (Butler and Wallingford, 2018; Ciruna et al., 2006; McGreevy et al., 2015; Nishimura et al., 2012; Ossipova et al., 2015b). In principle, this polarity may derive from the existing egg polarity, e. g. animal-vegetal molecular gradient, or forms in response to a localized cue. Our tissue manipulation experiments suggest that the neuroepithelial PCP is progressively specified along the anteroposterior body axis. We propose that PCP is not defined by a preexisting gradient but is induced by a planar signal from the dorsal blastopore lip.

## Results

### Progressive posterior-to-anterior acquisition of PCP in the *Xenopus* neural plate

In *Xenopus* midneurula embryos, endogenous Vangl2 and exogenous fluorescent Prickle3 form crescent-shaped aggregates at the anterior borders of neuroepithelial cells

(Suppl. Figure 1)(Ossipova et al., 2015b). The anterior position of these aggregates has been established based on the staining of cells mosaically-expressing exogenous fluorescent PCP proteins and in morpholino-mediated knockdowns (Chuykin et al., 2018; Ossipova et al., 2015b) and it has been verified in our experiments. To investigate the spatial and temporal dynamics of PCP proteins in the *Xenopus* neural plate (NP), we performed time-lapse imaging of embryos expressing HA-Vangl2 and GFP-Prickle3 (GFP-Pk3) from late gastrula to midneurula stages (Figure 1A-C’’, Supplementary Video 1). At stage 12.5, GFP-Pk3 aggregates were observed only in a few interspersed cells (Figure 1A-A’’). Shortly after, GFP-Pk3 crescents were visible in the posterior NP, while only a few small aggregates were evident in the anterior region (Figure 1B-B’’). GFP-Pk3-containing complexes became prominent in both the anterior and the posterior NP only by stage 15/16 (Figure 1C-C’’). Of note, the formation of GFP-Pk3 aggregates correlated with the areas of cell displacement that were visible during NP elongation (Suppl. Video 1). This analysis indicates that the NP acquires PCP in the posterior-to-anterior direction and in parallel with early morphogenetic events. This conclusion was confirmed by immunostaining for endogenous Vangl2. Vangl2 crescents were detectable only in the posterior NP at stage 13, but they became visible in the whole NP at stage 15/16 (Figure 1D-I). Together, these experiments show the progressive acquisition of neuroectodermal PCP.

**Figure 1.**
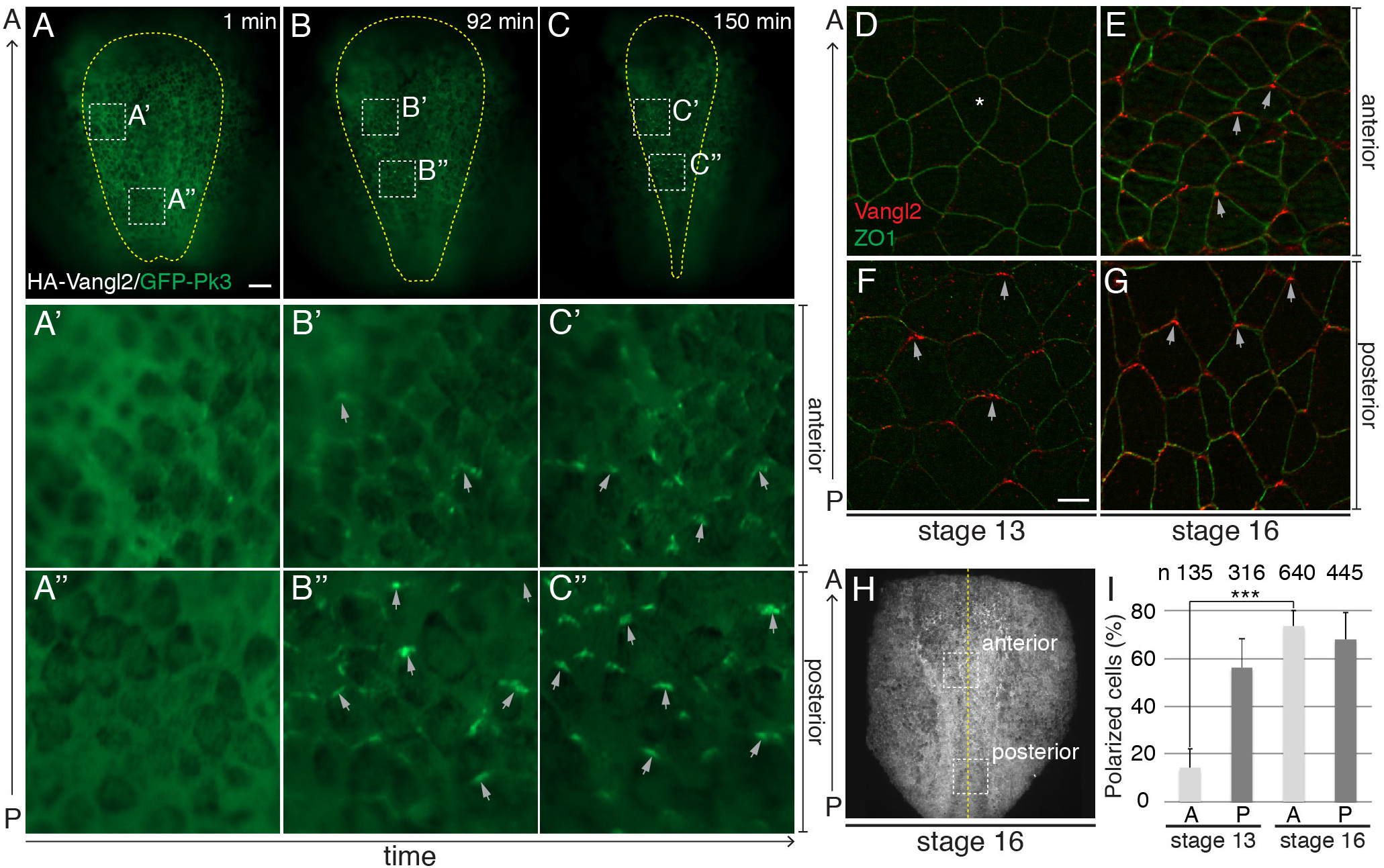
Progressive posterior-to-anterior acquisition of PCP in the *Xenopus* neural plate. (A-C’’) Still frames (dorsal view) of time-lapse imaging (stages 12.5 to 16) of a *Xenopus* embryo, expressing HA-Vangl2 and GFP-Pk3. Scale bar in (A), 100 μm, also refers to (B, C). Yellow line delimits the NP. Boxed regions in (A-C) are enlarged in A’-C”. Grey arrows, GFP-Pk3 crescent orientation. The results are representative of three independent experiments. (D-G) Vangl2 and ZO1 immunostaining of NP. ZO1 demarcates cell borders. Arrows, anterior Vangl2. Asterisk, lack of anterior Vangl2. The anteroposterior (A-P) axis is indicated. Scale bar, 10 μm, in (F), also refers to (D, E, G). (H) Stage 16 embryo at low magnification. Boxed regions, approximate positions of images in (D-G). Yellow line, midline. (I) Percentage of cells with polarized Vangl2 relative to the total number of ZO1-positive cells. n, number of cells per group. Means and s. d. are shown. Two-tailed Student’s *t* test, \*\*\**p* =1.5e^−6^.

### Posterior origin and planar propagation of the PCP cue

The posterior-to-anterior acquisition of PCP in the NP led us to hypothesize that the instructing cue originates at the posterior end of the embryo. Alternatively, PCP may independently develop in the anterior and posterior regions of the NP in a stage-dependent manner. To distinguish between these possibilities, we examined PCP in the NP, in which the continuity between the posterior and the anterior portions has been mechanically disrupted. A mediolaterally-oriented microsurgical incision (MLI) was introduced approximately in the middle of the neural *anlagen* at late gastrula stage (Figure 2A, Suppl. Figure 2A, B), and Vangl2 immunolocalization was analyzed in the operated and control embryos at stage 15. The incisions affected only the ectodermal layers, with the underlying mesendoderm remaining intact, thereby allowing to distinguish between ‘planar’ (*i. e.* transmitted in the plane) and ‘vertical’ (*i. e.* arising from the underlying mesendoderm) signals. Importantly, the microsurgical procedure did not compromise neural induction in manipulated embryos, as confirmed by the expression of the pan-neural marker Sox3 (Figure 2B-D). As expected, Vangl2 was enriched at the anterior borders of most cells of the control NP (Figure 2B-B”). In the embryos with the MLI carried out at stage 11.5, Vangl2 remained polarized posterior to the cut, but little if any polarization was observed anterior to the wound (Figure 2C-C’’, E). When the same surgery was done at stage 13, Vangl2 polarization was only slightly affected in the anterior NP (Figure 2D, E). Incisions made along the anteroposterior axis did not produce any significant change in Vangl2 crescent orientation (Suppl. Figure 2C-F’). These experiments indicate that the PCP signal requires tissue integrity for its transmission, propagates in the plane of the neuroectoderm, is distinct from neural inducers and acts at late gastrula-early neurula stages. These results also support our hypothesis regarding the posterior source of the PCP cue.

**Figure 2.**
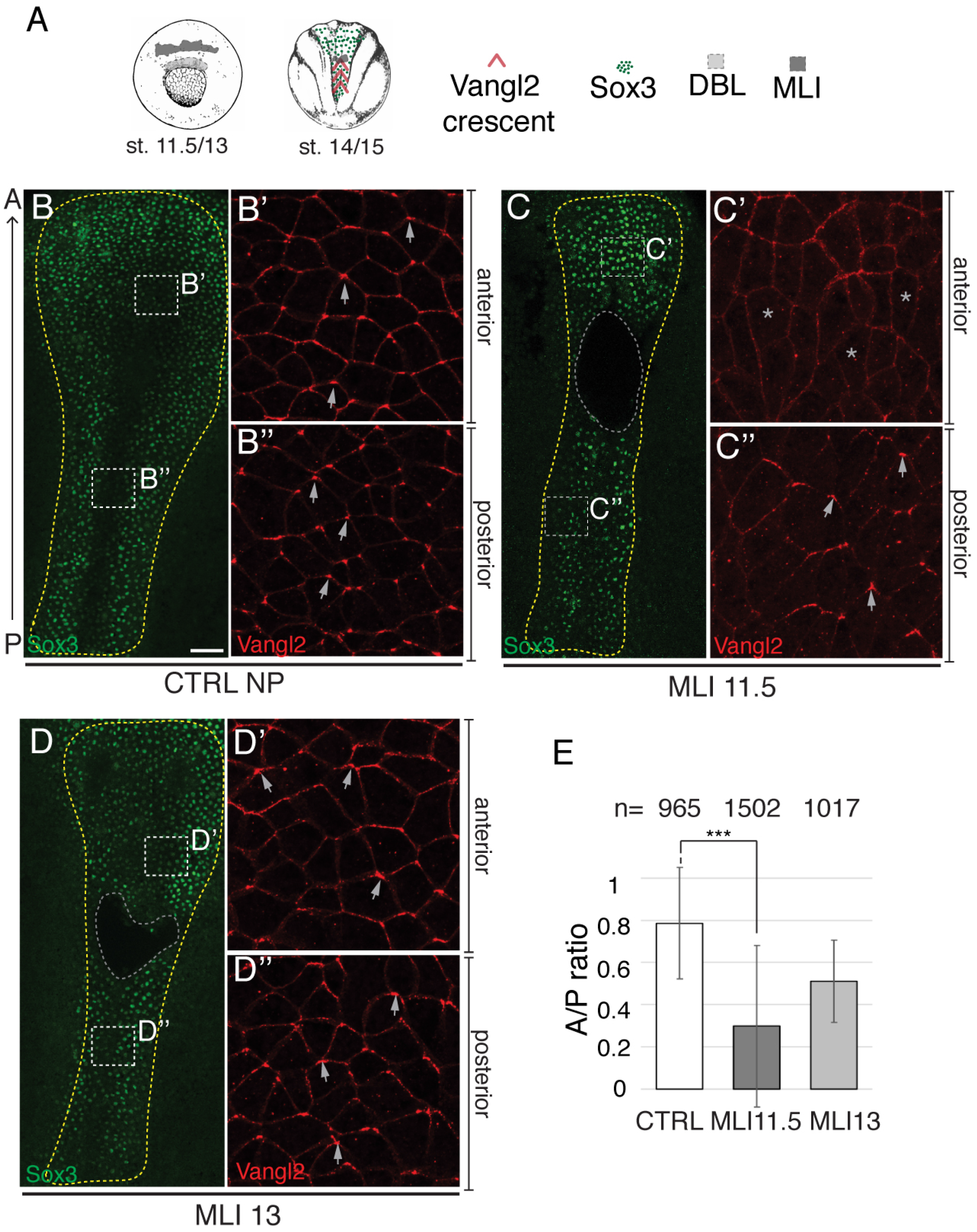
Posterior origin and planar propagation of the PCP cue. (A) Schematic of the experiment. Mediolateral incisions (MLI) were made at stage 11.5 (MLI11.5) or stage 13 (MLI13). (B-D’’) Representative neural plates (NP) double immunostained for Vangl2 and Sox3. (B-B’’) Control unmanipulated embryo (CTRL, stage 15, N=3). (C-C’’) MLI11.5 embryo (N=6). Asterisks in C’ mark the cells with lack of Vangl2 polarization. (D-D’’) MLI13 embryo (N=5). Boxed regions in (B, C, D) are magnified in (B’, B’’, C’, C’’ and D’, D’’). Yellow line delimits the NP. Grey line indicates MLI. Arrows show Vangl2 crescent orientation. N, number of embryos examined. Scale bar in (B), 100 μm, also refers to (C, D). (E) Ratio of the frequencies of polarized cells in the anterior and posterior NP (A/P ratio). n, number of cells per group. This is representative of 3-4 independent experiments. Means and s. d. are shown. Two-tailed Student’s *t* test, \*\*\**p*=0.00038.

To further address the origin and the transmission mode of the PCP signal, we analyzed Vangl2 distribution in exogastrulating embryos, in which the involution of mesendoderm does not occur (Holtfreter, 1933). Whereas in normal embryos, the neuroectoderm overlays the mesendoderm, in exogastrulae, these tissues are planarly connected by a stalk, corresponding to the posterior end of the embryo (Figure 3A, B). This tissue rearrangement allows to discriminate between planar and vertical signal transmission. In normal neurulae, both the NP and the gastrocoel roof plate (GRP), an equivalent of the mammalian node, are planar polarized (Antic et al., 2010; Borovina et al., 2010; Chu et al., 2016; Hashimoto et al., 2010). In exogastrulae, a change in the orientation of the NP and the GRP polarity vectors may provide insight about the source of the PCP cue (Figure 3C, D). Both the NP and GRP cells exhibited Vangl2 enrichment at anterior cell borders of stage 15 control embryos as expected (Figure 3E, E’). In the exogastrulae, Vangl2 accumulated in both tissues at the cell borders distal from the stalk (Figure 3F, F’), indicating that the polarizing signal emanates from the posterior region of the embryo. Moreover, since neuroectodermal PCP is preserved even after the displacement of the underlying mesendoderm, the signal is likely to propagate in the plane of the tissue.

**Figure 3.**
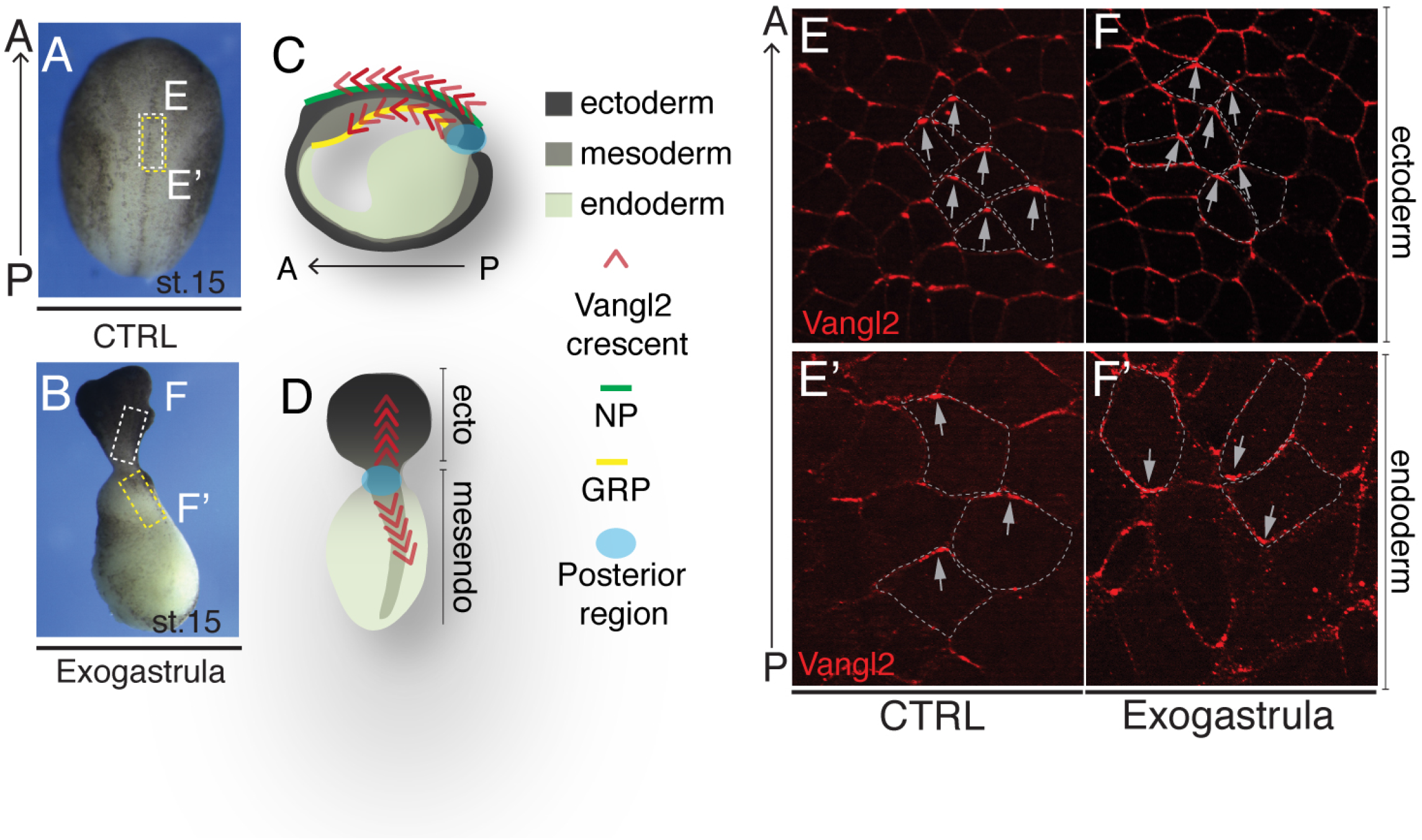
Vangl2 polarization in exogastrulae. (A, B) Brightfield images of control (CTRL, A) and exogastrula (B) embryos at neurula stage. (C, D) Schematic of normal (C) and exogastrula (D) embryos. (E-F’) Representative Vangl2 immunostaining of neuroectoderm (E, F, white boxes) or endoderm (gastrocoel roof plate (GRP) for normal embryos) (E’, F’, yellow boxes). Grey arrows, Vangl2 aggregate orientation. White dashes outline cell borders. Data represent three independent experiments with 2-6 embryos per group.

### The dorsal blastopore lip exhibits PCP-inducing activity

Our results suggest that the dorsal blastopore lip (DBL) at the posterior end of the embryo is the source of the PCP cue. To test this hypothesis, we grafted the DBL or the ventral epidermis (VE) of a donor embryo into the prospective NP of a recipient embryo at stage 12/12.5 and assessed Vangl2 localization at stage 15 (Figure 4A-C, Suppl. Figure 3). Vangl2 accumulation was clearly visible at the anterior cell borders in the entire NP of control embryos (Suppl. Figure 3C-E’). However, in recipient embryos, Vangl2 crescents oriented posteriorly in the cells within 150 μm of the DBL graft, yet remained unaffected at a distance (Figure 4D, F, Suppl. Figure 3F). PCP reversals were accompanied by the change in cell and tissue shape. Specifically, the tricellular junctions, frequently associated with the sites of Vangl2 accumulation (Ossipova et al., 2015b), were visible at the posterior cell edge rather than the anterior edge as in the control NP (Figure 4D, F, Suppl. Figure 4). Of note, we often observed radial orientation of Vangl2 crescents relative to the transplanted tissue (Suppl. Figure 4). By contrast, after VE grafting, Vangl2 aggregates retained the normal orientation at the anterior of each cell (Figure 4E, G, Suppl. Figure 3F), indicating that the observed PCP reversal was not due to the microsurgical procedure itself. Notably, ventral blastopore lip grafts also showed a weak PCP-inducing activity when grafted into the neural *anlagen* (Suppl. Figure 4).

**Figure 4.**
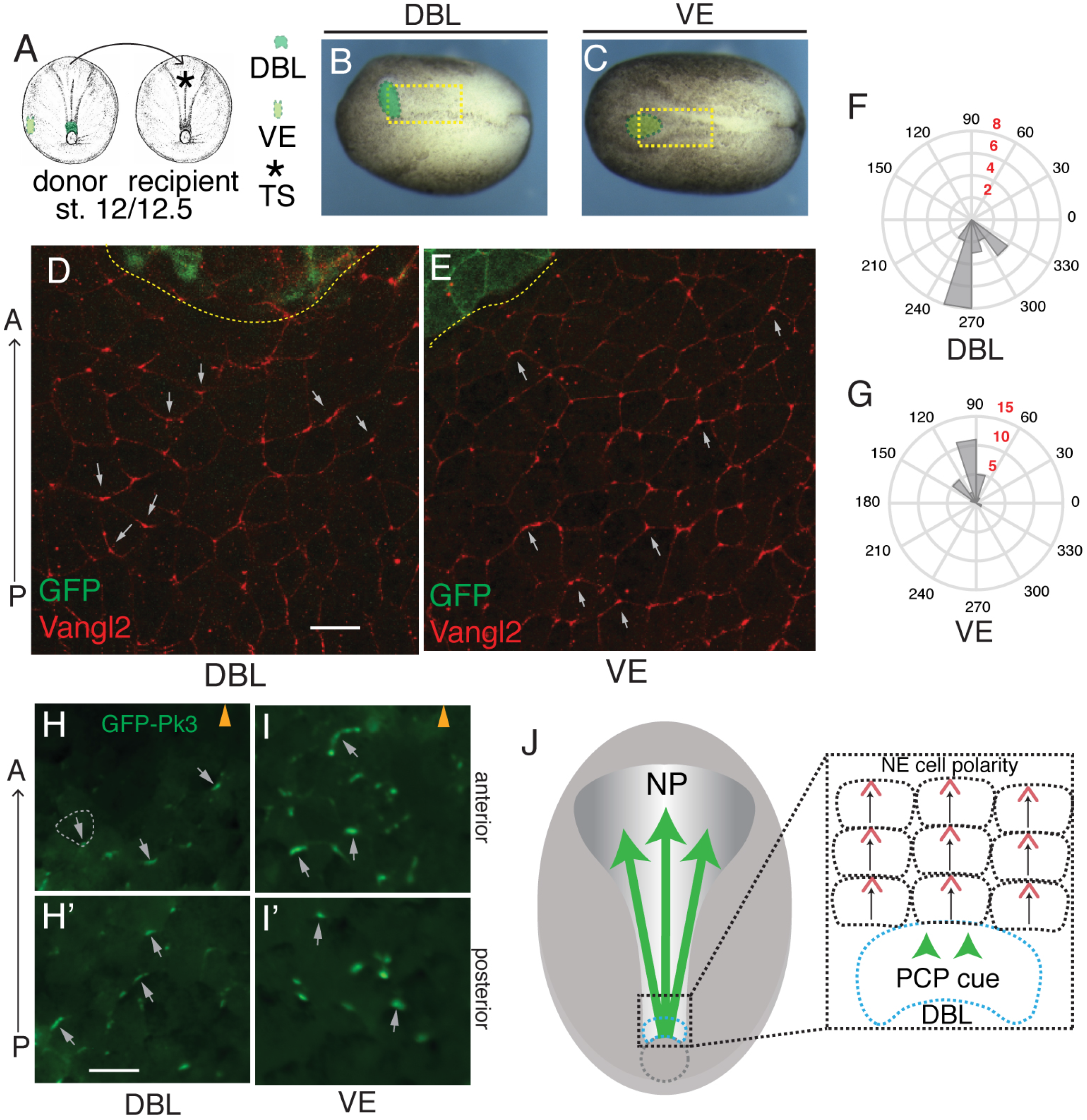
Dorsal blastopore lip grafts exhibit PCP-inducing activity. (A) Schematic of dorsal blastopore lip (DBL) and ventral fragment (VE) grafting experiment. TS, transplantation site (asterisk). (B, C) Brightfield images of neurula embryos grafted with DBL (B) and VE (C) (green) to the anterior NP. Yellow boxes approximate the regions shown in (D) and (E). (D, E) GFP and Vangl2 immunostaining of NPs grafted with DBL (D, n=7) or VE (E, n=3) at stage 12/12.5. Flag-GFP RNA is graft lineage tracer. Grey arrows indicate Vangl2 aggregate orientation. Scale bar, 70 μm, in (D), also refers to (E). (F, G) Rose plots quantify Vangl2 aggregate orientation within 150 μm from the graft. (F) DBL graft, (G) VE graft. Black numbers, vector angle. Red numbers, number of aggregates per bin. These data are representative of 4-5 independent experiments. (H-I’) Fluorescent images of NPs from DBL-graft recipient embryos (stage 15) expressing GFP-Pk3 and HA-Vangl2 (unlabeled). DBL (H, H’) or VE (I, I’) were grafted to the anterior NP at stage 12/12.5. Graft position is indicated by arrowheads in H and I. Images shown in H and I are proximal to the graft (within 150 μm) (labeled as anterior), whereas H’ and I’ are more distal, from the posterior region of the NP (labeled as posterior). White dashed line indicates cell borders in (H). Grey arrows show GFP-Pk3 crescent orientation. Scale bar, 60 μm, in (H), also refers to (H’, I, I’). The images are representative of 4-5 independent experiments, each including 2-6 embryos per group. (J) Planar induction of PCP during gastrulation. PCP is induced in the neuroectoderm by a signal from the dorsal lip and likely propagates across the tissue by a cell contact-mediated process.

To confirm these findings, we performed the grafting to recipient embryos expressing exogenous HA-Vangl2 and GFP-Pk3. Supporting the previous conclusion, DBL but not VE grafts caused GFP-Pk3 polarity reversals in the neuroectoderm in the majority of the recipient embryos (Figure 4H-I’). Together, these experiments demonstrate that the DBL is a source of the PCP instructing signal (Figure 4J).

## Discussion

This study evaluated the dynamics of PCP protein localization to investigate how PCP is established in the vertebrate NP. Our spatiotemporal analysis suggests that PCP is progressively established in the NP in the posterior-to-anterior direction. Also, the anterior NP does not acquire PCP, when ectoderm integrity is disrupted at the midgastrula stage. Based on this evidence, we propose that PCP is not derived from the oocyte polarity or instructed by a pre-established gradient, but is induced at the end of gastrulation by a signal or signals from the dorsal blastopore lip. Since only the ectoderm germ layer has been disrupted in our experiments, the PCP-instructing signal must be transmitted through the plane of the tissue. This conclusion is consistent with our findings that PCP is preserved in exogastrulae. By contrast, neural induction appears to only partly rely on planar signaling (Poznanski and Keller, 1997). We further found that blastopore lip grafts re-orient Vangl2 and GFP-Prickle3 crescents away from the graft. Taken together, these experiments argue that PCP is acquired in the neuroectoderm response to a planar cue originating in the DBL (Figure 4J). Thus, our work provides the first evidence of PCP induced by a dorsal lip transplant.

Whereas we have demonstrated that neuroectodermal PCP arises in response to inductive signaling, our current experiments do not reveal the identity of the PCP cue(s). The PCP-instructing signal could be a diffusible molecule or a mechanical force that is transmitted through the plane of the tissue. Furthermore, core protein bridges between neighboring cells are likely to contribute to PCP, reflecting domineering non-autonomy that was described in both *Drosophila* and vertebrate models (Mitchell et al., 2009; Sienknecht et al., 2011; Vinson and Adler, 1987). These possibilities are not mutually exclusive. Importantly, the PCP-instructing activity of the DBL appears distinct from the organizer signals mediating neural induction (De Robertis and Kuroda, 2004; Harland, 2000). First, mediolateral incisions prevent PCP establishment in the anterior NP, but do not interfere with neural induction as manifested by Sox3 expression. Second, both the DBL and the VBL contain PCP-instructing activity, even though the VBL is not able to induce neural tissue. Third, PCP reversals in the NP do not depend on graft orientation, arguing against an axis-inducing activity being responsible for PCP.

Evidence has been accumulating for biochemical signals and physical forces as alternative long-range PCP cues. Wnt gradients have been proposed to control PCP by modulating Frizzled (Minegishi et al., 2017; Wu et al., 2013) or Vangl2 activity (Gao et al., 2011; Yang et al., 2017), or by affecting tissue growth and morphogenesis (Sagner et al., 2012). In fact, our previous study indicated that Wnt11b is both necessary and sufficient for early PCP generation in Xenopus (Chu and Sokol, 2016), but whether this is a direct effect on the PCP signaling pathway remains unclear. Of note, Wnt ligands have been reported to regulate Myosin II via RhoA and ROCK activation (Habas et al., 2001; Marlow et al., 2002; Weiser et al., 2009). Importantly, Myosin II function may be essential for PCP (Ossipova et al., 2015b; Strutt et al., 1997; Winter et al., 2001). With Myosin II being a well-known force-producing molecule (Vicente-Manzanares et al., 2009), its activation by Wnt signaling would support the proposed role of mechanical strains in PCP (Aigouy et al., 2010; Aw et al., 2016; Chien et al., 2015). Whereas our experiments point to the existence of early signals that are responsible for *de novo* induction of PCP, future studies are needed to understand how this signaling is sensed and transmitted by core PCP components to connect biochemical to physical signals during PCP induction.

## Acknowledgements

We thank Dominique Alfandari for the anti-Sox3 antibody, Elena Torban and Robert Vignali for the comments on the manuscript, members of the Sokol laboratory for discussions. We acknowledge the MSSM Microscopy Core for the use of confocal microscopes and helpful advice. This work has been supported by the National Institutes of Health grants GM122492 and NS100759 to S. Y. S.

## Methods

### Xenopus embryos and microinjections

*Xenopus laevis* adults were purchased from Nasco and maintained according to the NIH Guidelines for the Care and Use of Laboratory Animals. Animal care and use were in accordance with the guidelines established by the Icahn School of Medicine at Mount Sinai. *In vitro* fertilization and embryo culture were performed as described previously (Dollar et al., 2005). Staging was according to (Nieuwkoop and Faber, 1967). Embryo microinjections were carried out at the 8-to-32-cell stage in 3 % Ficoll 400 (Pharmacia) in 0.5 X Marc’s modified Ringer’s (MMR) solution (Newport and Kirschner, 1982), 10 nl of RNA solution was injected into one or two blastomeres. Embryos were cultured in 0.1 X MMR either at 14°C or room temperature (RT). All experiments were repeated at least three times, with 3-10 embryos per group.

### Plasmids, mRNA synthesis and knockdown experiments

Capped RNAs were synthetized by *in vitro* transcription from the T7 or SP6 promoters using mMessage mMachine kit (Ambion) from the plasmids encoding HA-Vangl2 (Ossipova et al., 2015b), GFP-Pk3 (Chu et al., 2016), Flag-GFP (Chu et al., 2016). RNAs were injected at the following doses: HA-Vangl2 (60 pg), GFP-Pk3 (150 pg), Flag-GFP (100 pg), memRFP (100 pg).

### Embryo microinjections and microsurgical manipulations

For incision experiments, embryos were cultured in 0.1 X MMR until stage 11.5 or 13, then were devitellinized and an incision was made along the desired axis using a glass needle in 0.1 X MMR. The incisions were made through the ectoderm only, with mesendoderm remaining unaffected. The embryos developed in 0.1 X MMR at RT until midneurula stage, wound healing was prevented by repetitive separation of the wound margins with a glass needle every 10-20 min. Manipulated embryos and control siblings were fixed at stages 15-16 for immunofluorescence analysis.

For grafting experiments in wild type recipients, donor embryos were injected in one ventral or dorsal marginal blastomere with Flag-GFP RNA (100 pg, lineage tracer). Injections were performed at the 32-cell stage to minimize gastrulation defects and assure transplantation of the desired region. For grafting experiment in recipient embryos expressing exogenous PCP markers, recipient embryos were coinjected with HA-Vangl2 (60 pg) and GFP-Pk3 (150 pg) RNA in two animal dorsal blastomeres at the 8-cell stage, donor embryos were injected with memRFP RNA (100 pg) in one animal ventral or dorsal marginal blastomere at the 32-cell stage. Both donor and recipient embryos were cultured in 0.1 X MMR until stage 12/12.5, then transferred in 1 X MMR with 50 μg/ml of gentamicin (Sigma) for the microsurgical procedure. Both recipient and donor embryos were devitellinized, the DBL, VBL or VE (corresponding to an area of approximatively 0.02/0.03 mm^2^) were excised from the donor embryo, the explanted tissue comprises all germ layers. An incision was then made in the neural *anlagen* of the recipient embryo, in the medial-anterior position, and the graft was transferred to the recipient embryo. The superficial layer of the graft remained on the surface of the embryo, but it was randomly oriented with respect to the recipient’s body axis. Grafted embryos were cultured in 1 X MMR for about 15-30 min, until wound healing was completed and then transferred to 0.1 X MMR, cultured until stage 14/15 at RT and fixed for staining and imaging.

Exogastrulation was induced by placing embryos into 1 X MMR at stage 7, control sibling embryos were cultured in parallel in 0.1 X MMR. The exogastrulae and the control embryos were devitellinized at stage 10 and cultured at 14°C until stage 14/15, at which point they were fixed for immunofluorescence analysis.

### Tissue preparation, immunostaining and live imaging

For fluorescence imaging of exogenous GFP-Pk3, RNA-injected embryos were devitellinized and fixed in MEMFA (0.1 M MOPS pH 7.4, 2 mM EGTA, 1 mM MgSO_4_ and 3.7 % formaldehyde)(Harland, 1991), for 30-60 min at RT, washed in 1 X phosphate buffered saline (PBS) (5 min at RT) and stored in 1 X PBS at 4°C. Embryos were fixed at 11.5 or 15/16 in specific experiments, as described above. NPs or ventral ectoderm were manually dissected from fixed embryos with a scalpel in 1 X PBS, and mounted on slides using Vectashield mounting medium (Vector).

For immunostaining, embryos were devitellinized, fixed in 2 % trichloracetic acid (TCA) in water for 30-60 min and stained as described (Ossipova et al., 2015b). Fixed embryos were rinsed and stored in 1 X PBS at 4°C for a maximum of 5 days. NPs were manually dissected from fixed embryos, washed in 1 X PBS with 0.3 % Triton X100 for 2 hrs at RT, then incubated in primary blocking solution (5 % normal donkey serum, 1 % BSA, 1 % DMSO, in 1 X PBS with 0.1 % Triton X100) for 2 hrs at RT. Samples were incubated with primary antibodies diluted in the blocking solution overnight (O/N) at 4°C, then washed 5 times, 1 hr each, in 1 X PBS with 0.1 % Triton X100, then incubated with secondary antibodies diluted in secondary blocking solution (0.1 % BSA, 1% DMSO in 1 X PBS with 0.1 % Triton X100) O/N at 4°C. After 5 washes of 1 hr each in 1 X PBS with 0.1 % Triton X100, NPs were mounted for imaging as described above.

The following antibodies were used: rabbit anti-Vangl2, 1:200 (Ossipova et al., 2015a), mouse anti-Sox3, 1:25 (hybridoma clone 5H6, a gift of Dominique Alfandari), mouse anti-GFP (B2, Santa Cruz), mouse anti-ZO1, 1:200 (Invitrogen), Cy3-conjugated donkey anti-rabbit IgG, 1:250 (Jackson ImmunoResearch), Cy2-conjugated donkey anti-mouse IgG, 1:250 (Jackson ImmunoResearch), Alexa Fluor488-conjugated donkey anti-mouse IgG, 1:250 (Jackson ImmunoResearch).

Fluorescent images were captured using AxioImager microscope (Zeiss) and Axiovision imaging software, or LSM880 Airyscan confocal microscope (Zeiss) and Zen (black edition) imaging software. Z-stacks were acquired and maximum intensity projections were obtained by processing imaging files with Fiji (ImageJ).

For live imaging, embryos were injected with HA-Vangl2 (60 pg) and GFP-Pk3 (150 pg) RNAs in both animal dorsal blastomeres at 8-cell stage and let develop until stage 11.5. Then the embryos were mounted in 0.8 % low melting point agarose in 0.1 X MMR in an imaging chamber formed by a glass slide and a glass coverslip separated by a spacer. Embryos were imaged from the dorsal side, from late gastrula to midneurula stages, one frame taken every 2 min. Three to five embryos were imaged in each experiment. AxioZoom fluorescent microscope (Zeiss) and Zen (blue edition) imaging software were used.

### Quantification and statistical analysis

Vangl2 crescent scoring in the MLI experiments was performed using Fiji “analyze particles” command (ImageJ). The ratio between the frequencies of polarized cells in the anterior and posterior NP was calculated by the following formula: A/P ratio= (N_CA_/tN_A_) / (N_CP_/tN_P_) (N_C_, number of crescents; tN, total number of cells; A, anterior; P, posterior). For the analysis of Vangl2 localization in wild type embryos at different developmental stages, the frequency of cells showing Vangl2 anterior localization was calculated over the total number of neuroectodermal cells expressing ZO-1. Average and s.d. were calculated with Microsoft Excel. Histograms representing data quantifications were obtained using Microsoft Excel.

Vangl2/GFP-Pk3 aggregate orientation was quantified using Fiji (ImageJ). For each sample, all cells with clear crescents within an area of 0.25 mm^2^ posterior to the grafted tissue were analyzed. For each cell, a line connecting the two extremities of the crescent and an arrow perpendicular to this line were drawn, and the angle (α) between this arrow and the antero-posterior axis was measured, as shown in Suppl. Figure 1A’, B’. Anterior, lateral and posterior orientation of the crescents were defined by the α angle as described in Suppl. Figure 1. For each cell, the distance from the graft was also measured. For the control NP, the anterior and posterior regions were defined as the regions within 250 μm from the geometric center in the respective directions. Due to the size and the position of the transplant, the analyzed regions were subdivided into anterior (within 150 μm from the graft) and posterior (further than 150 μm from the graft) as shown in Suppl. Figure 3A. Measured crescent angles were presented as rose plots obtained by using Matlab_R2019a and also shown as function of their distance from the graft in scatter plots obtained using Microsoft Excel.

For linear statistic mean comparison, Student’s t-test was used. Circular statistics analysis was performed with Matlab_R2019a Circular Statistic Toolbox. The circular means and circular variances were calculated for each analyzed sample and plotted into bubble plots using Microsoft Excel. In each experiment, pairs of samples were compared. Pair-wise comparisons have been performed in each case, and the percent of comparisons with p<0.05 is shown.

---

**Suppl. Video 1. Propagation of PCP in the posterior-to-anterior direction in *Xenopus* neural plate.**

Time-lapse imaging of polarized GFP-Pk3 aggregate formation during neurulation. *Xenopus* embryos expressing HA-Vangl2 and GFP-Pk3 RNAs. Dorsal view is shown. Note progressive posterior-to-anterior formation of GFP-Pk3 crescents. The movie is representative of three independent experiments.

**Suppl. Figure 1.**
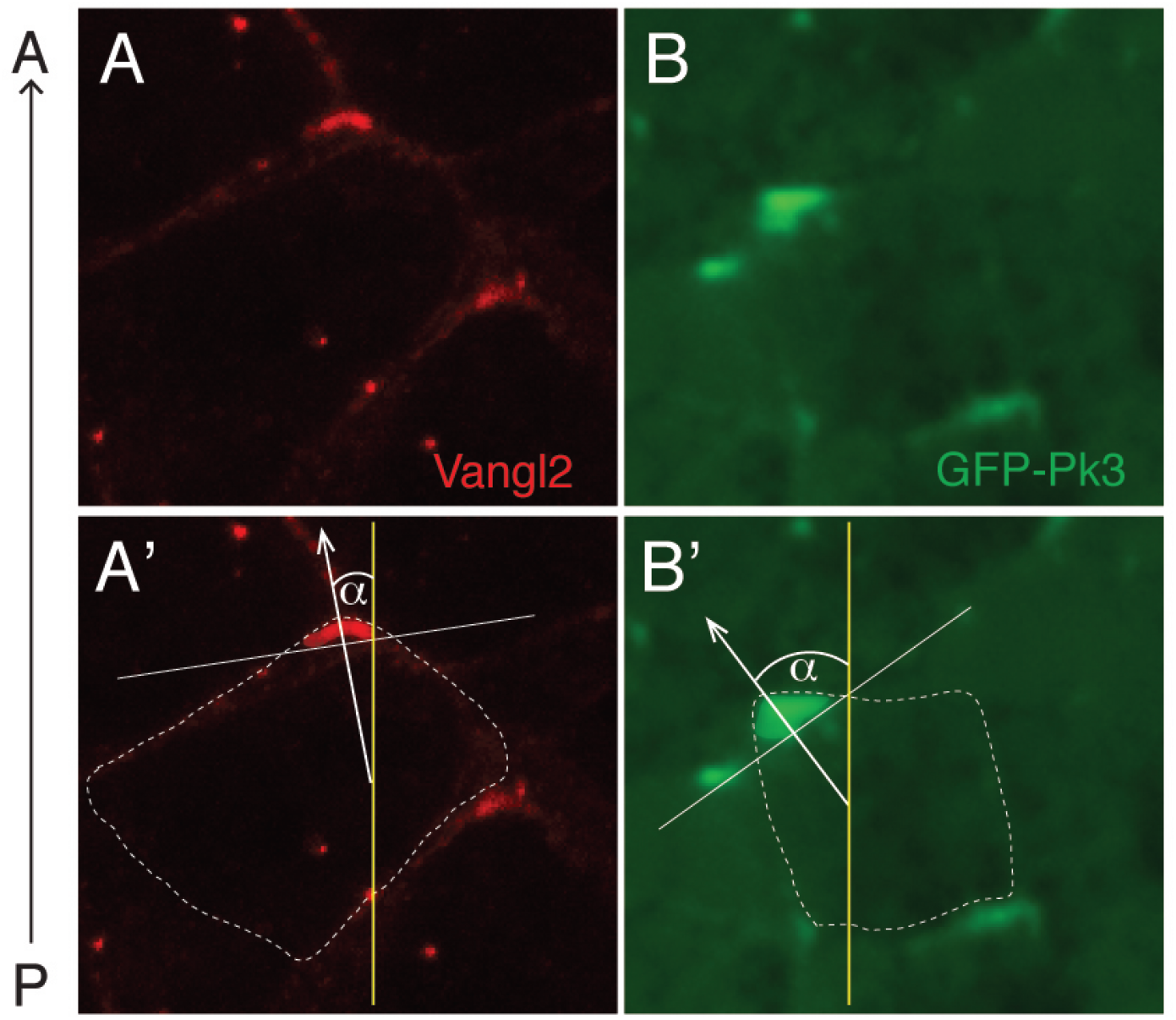
Scoring PCP aggregate orientation in the neural plate. (A, B) Neuroepithelial cells exhibit anteriorly polarized endogenous Vangl2 (immunofluorescence) (A), or exogenous GFP-Pk3 (GFP fluorescence) in the presence of exogenous Vangl2 (unlabeled) (B). (A’, B’) Scoring method for the angle (α) that represents the orientation of the polarity vector (white arrow) for Vangl2 (A’) and GFP-Pk3 (B’) aggregates.

**Suppl. Figure 2.**
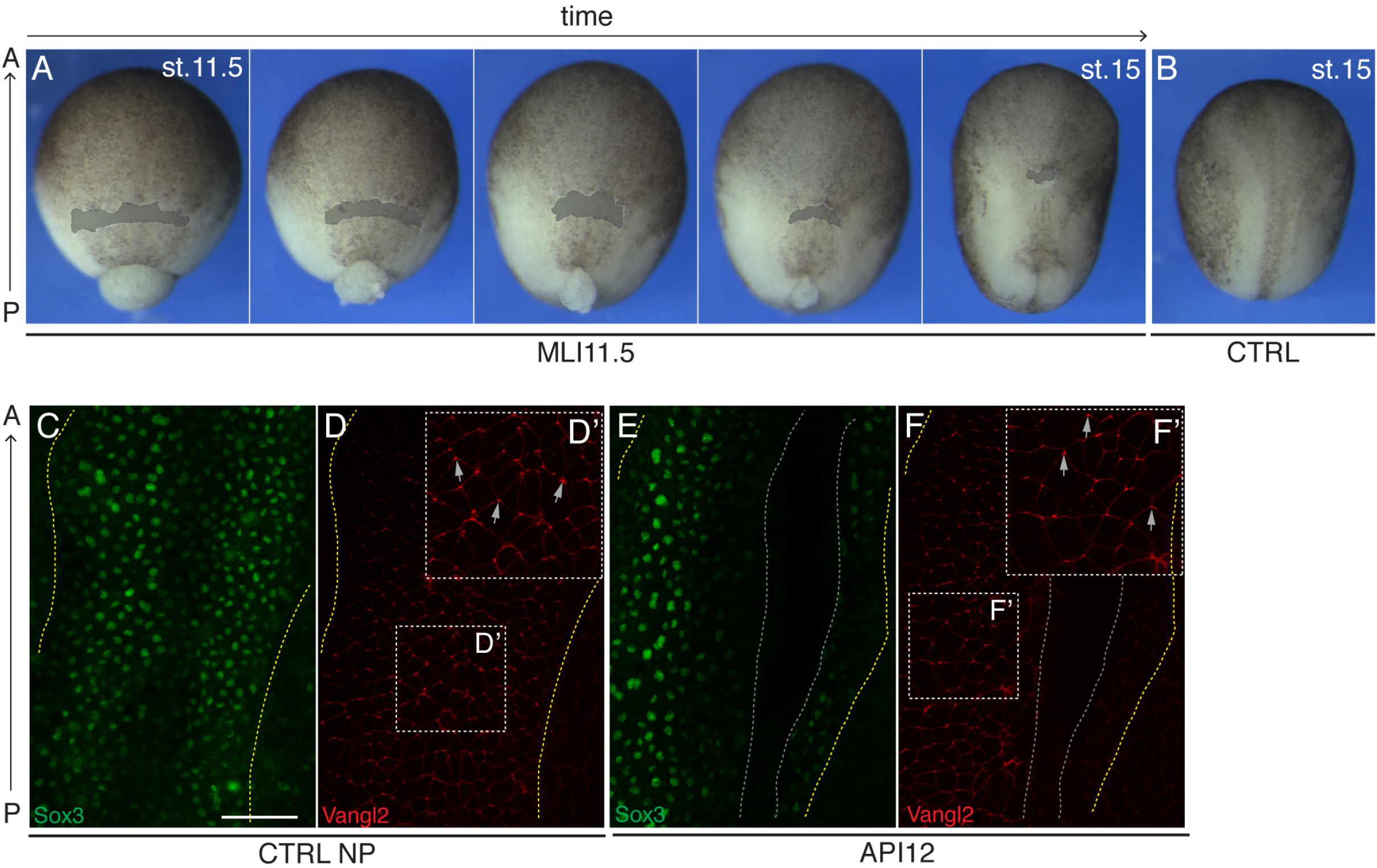
Embryo morphology and Vangl2 polarization after microsurgical incisions. (A) Brightfield timecourse images of an embryo (stages 11.5-15) with mediolateral incision (MLI) made at stage 11.5. Note the changes in embryo and wound shape. (B) Control embryo at stage 15. (C-F’) Sox3 and Vangl2 immunostaining of stage 15 control embryo (C, D) and embryo with anteroposterior incision (API12) (E, F) made at stage 11.5/12. Boxed regions in (D, F) are magnified in (D’, F’). Yellow lines delimit the neural plate, grey arrows show Vangl2 crescent orientation. Grey lines in (E, F) indicate the incision. The data are representative of three independent experiments with 3-6 embryos per group.

**Suppl. Figure 3.**
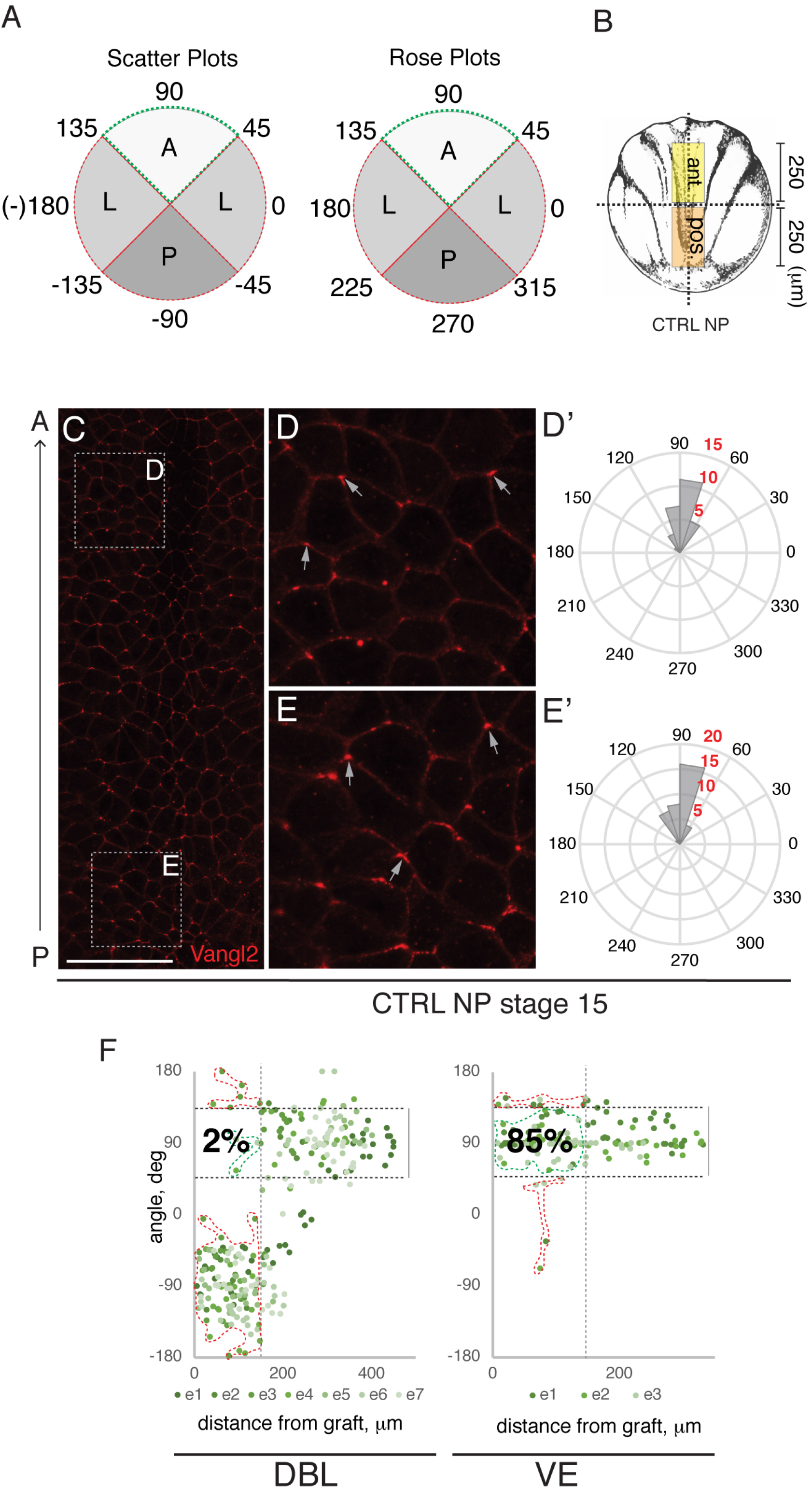
Quantification of Vangl2 aggregate orientation in the neural plate. (A) Schematic representation of the methods used: scatter plots and rose plots (see Methods for detailed description). (B) Scheme of the neural plate with indicated areas of scored Vangl2 orientation. (C-E’) Quantitation of Vangl2 polarization in the neural plate (NP) at stage 15. (C) Vangl2 immunostaining of a control NP at low magnification. Boxed areas in correspond to the anterior and posterior regions magnified in D and E, respectively. Grey arrows indicate Vangl2 crescent orientation. Scale bar in C, 100 μm. (D’, E’) Rose plots showing Vangl2 aggregate orientation. (F) Scatter plots of Vangl2 polarization in embryos grafted with DBL and VE. PCP vector is shown as function of the distance from the transplant. Shades of green represent different embryos. Green dots between black dashed lines represent cells with anteriorly localized Vangl2 crescents. Dashed shapes indicate normal (green) or altered (red) orientation. Grey dashed vertical line marks 150 μm from the graft. Percentages refer to the number of cells with normal anterior orientation within 150 μm from the graft.

**Suppl. Figure 4.**
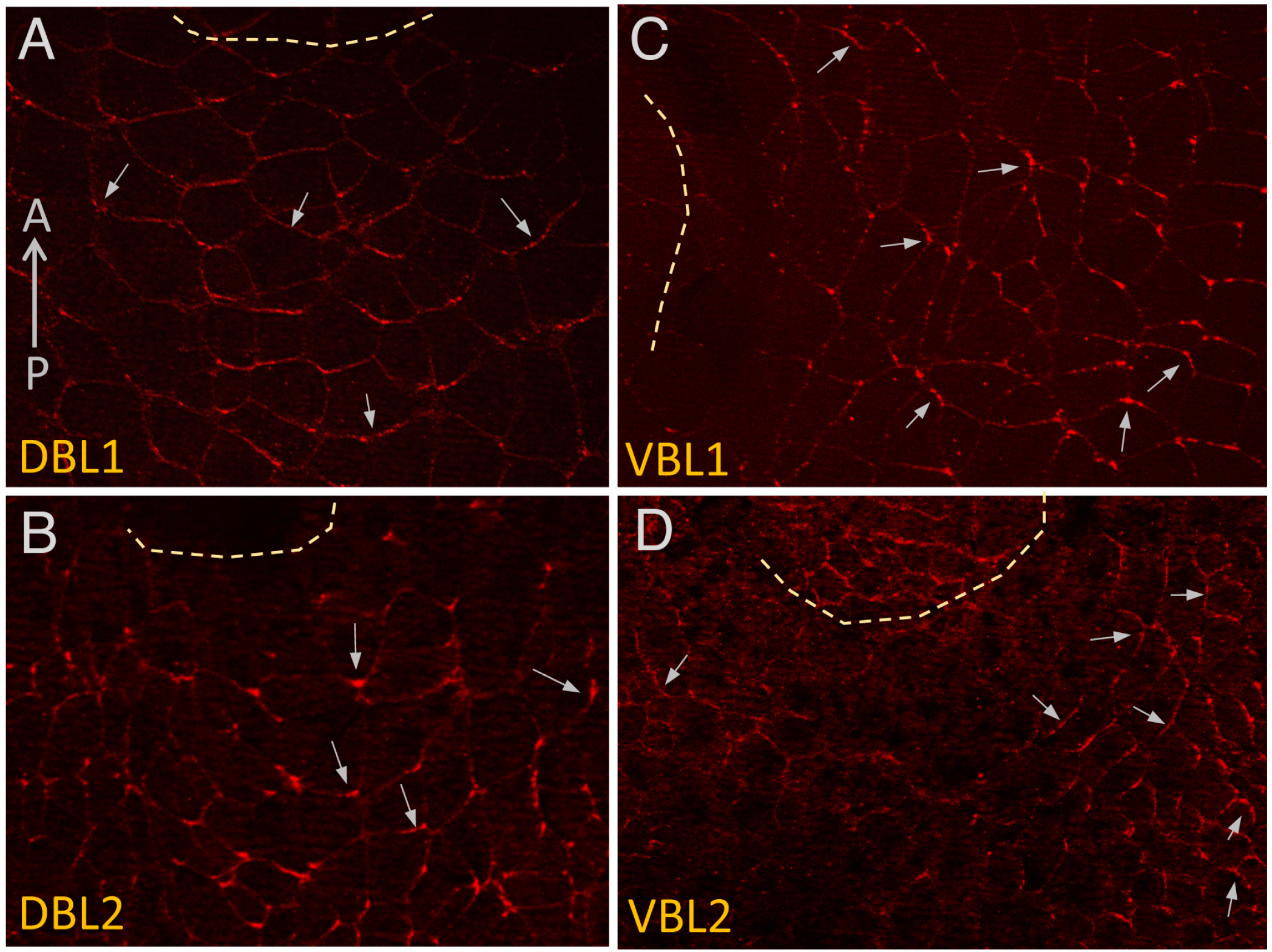
Both dorsal and ventral blastopore lip grafts cause PCP reversals in recipient neural plates. (A-D) Vangl2 crescent orientation in the proximity of DBL (A, B) and VBL (C, D) grafts. Grafting was done as described in Figure 4. Two NP immunostained for Vangl2 are shown for each type of graft, which are representative of 4-7 independent experiments. Grey arrows indicate individual cell polarity based on Vangl accumulation. Approximate graft positions (dashed lines) and the orientation of the anteroposterior (AP) axis (refers to all panels) are shown.

